# Mapping nanoscale forces and potentials in live cells with microsecond 3D single-particle tracking

**DOI:** 10.1101/2022.06.27.497788

**Authors:** Shangguo Hou, Chen Zhang, Anastasia Niver, Kevin Welsher

## Abstract

3D single-particle tracking has the potential to resolve the molecular level forces which dictate particle motion in biological systems. However, the information gleaned from 3D single-particle tracking often cannot resolve underlying nanoscale potentials due to limited spatiotemporal resolution. To this end, we introduce an active-feedback 3D tracking microscope that utilizes silver nanoparticles (AgNPs) as probes to study intricate biophysical events in live cells at the nanometer and microsecond scales. Due to this extremely high and durable scattering photon flux of the plasmonic particles, 1 MHz sampling frequency at nanometer precision in all three dimensions can be achieved over an unlimited observation times. In this work, we applied microsecond-sampling, active-feedback 3D single-particle tracking to investigate the interaction between AgNPs and nanoscale filopodium on the live-cell surface. The nanometer precision and microsecond sampling revealed that TAT peptide modified particles visit and dwell at local “hot spots” on the filopodium surface. The high sampling rate further enabled the calculation of the local forces and potentials within these nanoscale hotspots on the cylindrical surface of live cell filopodia. This study presents a promising tool to investigate intracellular biophysical events with unprecedented spatiotemporal resolution and a pipeline to study nanoscale potentials on three-dimensional cellular structures.

## Introduction

Tracking particles in three dimensions can reveal the molecular forces and potentials that act on a moving particle in a complex environment (*1*). In short, a trajectory of a single particle is a map of its interactions with the environment. For example, a particle may randomly diffuse among areas with different viscosity, be pushed or pulled by molecular motors, be physically trapped in a confined region due to steric obstruction, or be tethered in a potential well by molecular interactions. However, the information gleaned from these trajectories is strongly determined by the spatiotemporal precision of each point in the path. Interactions that are smaller than the localization precision or more fleeting than the temporal resolution will go unresolved. Furthermore, the trajectories must be long enough to allow the particle to map out an entire surface and measure its diffusive and other properties with sufficient resolution. Pulling the molecular scale information out of single trajectories requires spatiotemporal precision commensurate with forces and potentials at play.

Traditional 3D single-particle tracking methods have shown that particles will undergo random (Brownian) motion in solution, as well as a variety of confined diffusions with distinct geometries, such as diffusion along linear axes (*2*), cell surfaces (*3*), or isolated compartments (*4*). Decoding the hidden information behind diffusion requires high spatiotemporal resolution. An example is the progressive and retrogressive stepwise motion of motor proteins, which exhibit a step size of 8 nm (*5*). Resolving such features requires methods with localization precision on the order of several nanometers (*6*). Recently, due to the advances in localization precision, it has been revealed that previously considered non-moving particles undergo restricted diffusion, and that the difference in the slow diffusive speed correlates with the fate of such vesicles (*7*). Despite stunning progress that has been made in localization precision over the past two decades, advances in temporal resolution have fallen behind, largely due to the limited emission capacity of conventional fluorophores. This speed limit is magnified in 3D localization, which leaves an underexplored field to be fulfilled with methodologies with high spatiotemporal resolution. A method with several nanometers localization precisions, sub millisecond temporal resolution, and large field of view in 3D is needed to extract maximal information from particle trajectories.

One candidate for such a method is real-time 3D single-particle tracking (RT-3D-SPT) (*8-16*). RT-3D-SPT utilizes active-feedback mechanisms to lock the target in the microscope focus, enabling long duration and high spatiotemporal resolution observation in three dimensions. RT-3D-SPT is emerging as a powerful tool for the study of biological interactions due to its ability to unveil heterogeneity hidden within ensemble-averaged measurements, as well as its compatibility with living systems (*17*). However, there are several limitations to current RT-3D-SPT methods. First, conventional fluorescent particles widely used in RT-3D-SPT yield only a limited number of photons before undergoing irreversible photobleaching, limiting the total duration of single particle trajectories. Second, the emission rate of these particles is also limited acts as a barrier to the temporal resolution of RT-3D-SPT, which is limited only by photon collection rate. Recently, artificial scattering particles have gained substantial attention due to their superior photostability and biocompatibility. Various studies (*18-20*) have demonstrated the ability to obtain a high photon flux and high precision from using scattering particles such as gold or silver nanoparticles (NPs) as the probe in single-particle microscopic studies. However, most common scattering particle detection methods, such as dark-field microscopy and 3D-iSCAT, are limited by the lack of axial resolution (*21, 22*) or imaging depth (*23*), which prevent us from exploring the further benefit of scattering particles with almost unlimited photon fluxes in biological imaging. On the other hand, RT-3D-SPT, which has large imaging depth, single-photon sensitivity and up to tens of MHz photon count capability, promises to be an idea platform for the scattering observation in biological studies. While several studies have demonstrated the capability of tracking highly scattering NPs with RT-3D-SPT (*24-26*), these methods have thus far only been demonstrated in vitro and have not been translated to a live-cell setting.

Plasmonic resonance on metal surfaces has been widely used in single-molecule sensing, Raman scattering detection, and super-resolution imaging (*27-33*). Unlike fluorescence, surface plasmon resonance scattering displays distinct advantages. First, since the size of the NPs (100 nm or less) is much smaller than the excitation laser wavelength, the resulting plasmonic resonance will occur among electrons oscillating in a specific, directional axis and therefore resulting in polarized light even if the incident beam is not polarized. Therefore, the scattered light can be removed from the incident and reflected light using a polarizer. Moreover, as the scattering light has the same wavelength of incident light, background from autofluorescence can be removed with a narrow bandpass optical filter (Fig. 1a). What’s more, the scattered photons of the NPs are also not susceptible to photobleaching, quenching, or photoblinking. These features make scattering NPs ideal materials for real-time observation of cellular events with prolonged duration and high spatiotemporal precision.

**Figure 1.**
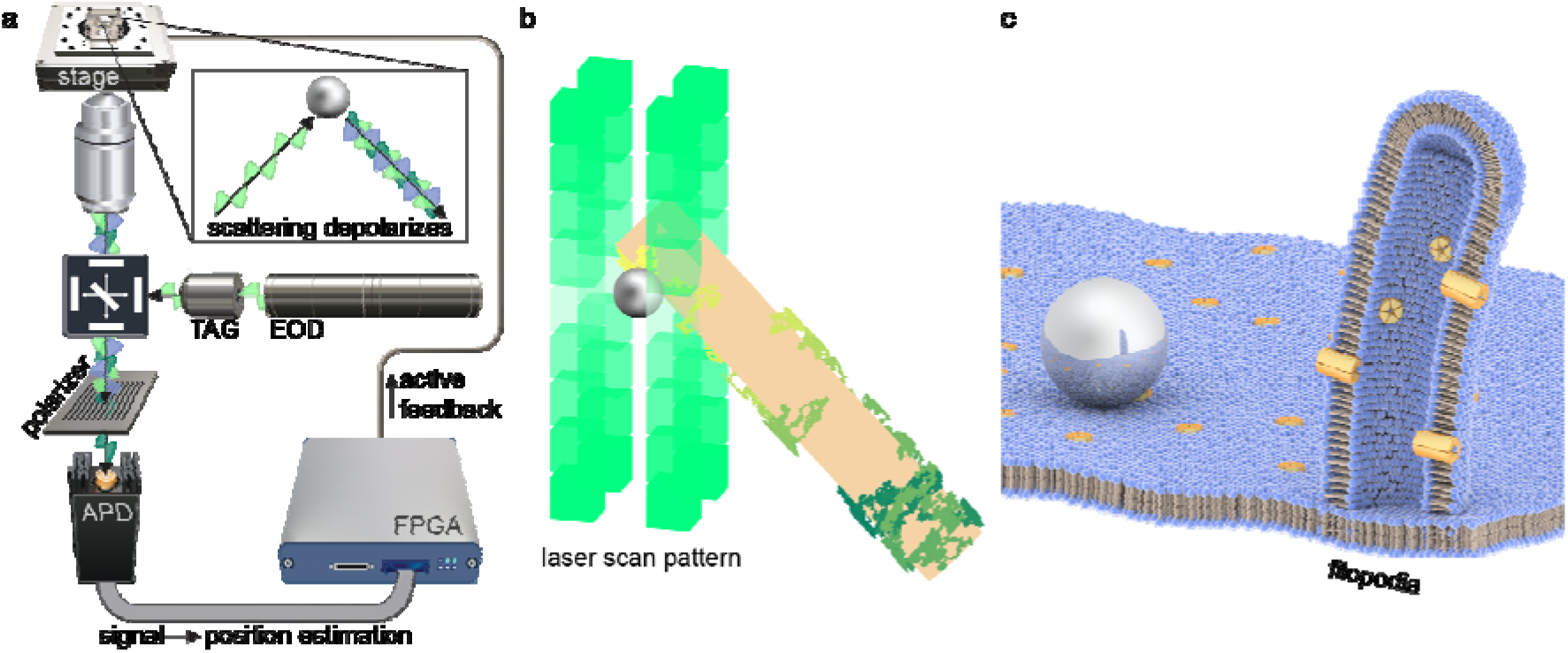
(**a**) Schematic of 3D-SMARTER. (**b**) Information-efficient sampling pattern implemented in 3D-SMARTER. (**c**) Interaction between AgNP and live-cell filopodia probed by 3D-SMARTER in this study.

In this study, we incorporated the detection of scattered and depolarized light from silver nanoparticles (AgNPs) into the previously reported 3D single-molecule active real-time tracking (3D-SMART) microscopy (Fig. S1) (*34*) with an information-efficient sampling pattern (*35*) (Fig. 1b, Fig. S2). This new method, incorporating high photon count rates and information-optimized scanning, is named 3D-SMARTER (3D-SMART with Enhanced Resolution). The combination of 3D-SMART and AgNPs made it possible to detect faster dynamics and finer cellular topological structures that cannot be revealed when the high spatiotemporal precision of RT-3D-SPT or the high and durable emitting capacity of metal particles alone. The detected depolarized scattering signal from AgNPs can be as high as 3 million per second, bringing unprecedented spatial and temporal resolution over three dimensions, while maintaining a large tracking depth and long observation time. In this work, we obtained 2.5 ± 0.2 nm localization precision in XY and 4.3 ± 0.5 nm precision in Z over a 50 μm axial range, with a temporal resolution down to 1 microsecond. The high spatial and temporal resolution of 3D-SMARTER enables observation of step motion of motor proteins in live cells and particle diffusion on complex, nanoscale 3D structures on the cell surface (Fig. 1c). Not only did the high spatial resolution of 3D-SMARTER carve out a super-resolved structure of cell surface filopodia, the high temporal resolution uncovered local hot spots on the surface of these nanoscale cylindrical protrusions that were repeatedly visited by single particles. Furthermore, the rapid sampling enabled mapping of force vectors and potentials within these hot spots, untangling them from random Brownian motion and localization uncertainty. These analyses are only possible with spatial and temporal precision better than the feature in question. 3D-SMARTER is a powerful tool to study biological events with unprecedented spatiotemporal precision.

## Results

### Tracking of AgNPs with high precision deep within samples

3D-SMART has been previously applied to track fluorescent protein tagged virus-like particles in live cells (*36*) and single fluorescent Atto 647N dye labeled proteins and nucleic acids in solution (*34, 37*), but in each case the photobleaching of fluorescent tags limits the signal rate and observation time. 3D-SMARTER was built upon the previously reported 3D-SMART platform. Briefly, an electro-optic deflector (EOD) and tunable acoustic gradient (TAG) lens are applied to rapidly shift the laser focus in a restricted volume in the vicinity of the particle (1 μm × 1 μm × 2 μm, XYZ). Real-time photon arrival information is then used to estimate the particle position within the scan area and used to guide a piezoelectric stage to actively track the target. Two major changes were applied to utilize the exceptional brightness and unique scattering mechanism of AgNPs. First, a polarizer was added to the detection path. The polarization angle of this polarizer was set perpendicular to the polarization angle of the excitation laser to maximize contrast of the scattering nanoparticle (Fig. S3). Second, rather than the previously reported 5×5 grid Knight’s Tour pattern, an information-efficient 4-pixel pattern (4-Corners pattern) that only samples areas with high Fisher information was used (*35*). The information-optimized 4-Corners pattern makes it possible to extract maximal information from the scattering signal. Critically, with the same 20 μs pixel dwell time used, the 4-Corners pattern leads to 80 μs period of single scan cycle, dramatically shorter than the 500 μs cycle of the previously used Knight’s Tour pattern, enabling greatly enhanced sampling rate.

To evaluate the robustness of the 3D-SMARTER mechanism, 100 nm PEG-AgNPs (2 pM, KJW1912, NanoComposix) were diluted into water and tracked in aqueous solution. 3D-SMARTER was able to track freely diffusing AgNPs in aqueous solution with diffusive speeds up to 4 μm^2^/s (Fig. S4), comparable to previously reported results obtained from fluorescent materials with 3D-SMART. To evaluate the localization precision of the method, citrate-stabilized AgNPs (2pM, AGCB100-1M, NanoComposix) were diluted into PBS and particles bound to the coverslip were tracked with 3D-SMARTER. The intensity trace of an immobilized AgNP being actively tracked is shown in Fig. 2a. The depolarized scattering signal is extremely stable with a count rate of ∼3.7 MHz. This ample photon flux made it possible to achieve nanoscale precision in all three dimensions. The tracking precision obtained for immobilized AgNPs was 2.5 ± 0.2 nm, 2.4 ± 0.2 nm and 4.3 ± 0.5 nm, for X, Y, and Z, respectively (Fig. 2b). This high precision was maintained at significant depth within the sample. To demonstrate, 100 nm AgNPs were diluted to ∼1 pM into Fluoromount mounting medium (Sigma, F4680). The resulting suspension was sandwiched between two coverslips, heated to fix the sample, and mounted on the microscope to track immobilized AgNPs at various depths. As shown in Fig. 2c, nanoscale localization of the AgNPs could be achieved over a range of larger than 50 μm in Z, far beyond several micron imaging depth of other scattering-based detection methods (*38*). Importantly, the XYZ tracking precision did not vary with depth within the sample (Fig. 2d)

**Figure 2.**
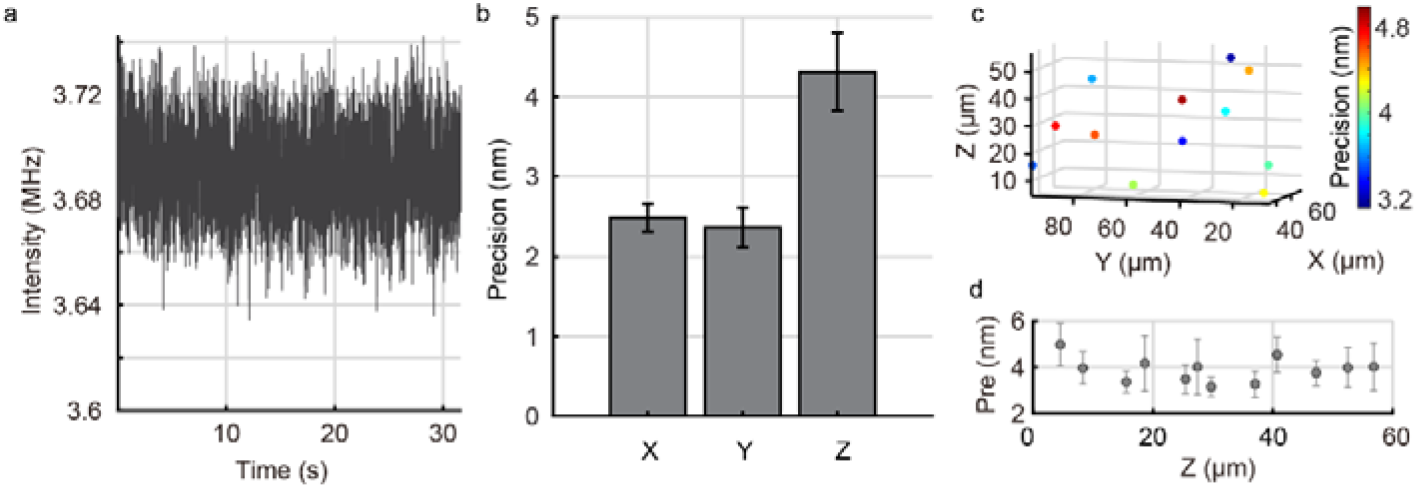
The precision and imaging depth of 3D-SMARTER. (**a**) Intensity trace from tracking an immobilized AgNP. Bin time: 1 ms. (**b**) Standard deviation of stage readout of X, Y, and Z axes while tracking of immobilized AgNPs. X, Y and Z precision was 2.5 ± 0.2 nm, 2.4 ± 0.2 nm and 4.3 ± 0.5 nm, respectively. Stage positions were sampled at 1 kHz. (**c**) Positions of 12 different AgNPs immobilized in Fluoromount medium at different depths in Z. (d) Tracking precision as a function of tracking depth. The coverslip surface is located at Z = 0 μm.

### Characterization of cargo motion in live cells

To demonstrate the benefits of the nanometer and millisecond precision of 3D-SMARTER, we applied 3D-SMARTER to quantify the steps of intracellular trafficking via motor proteins. The motion of motor proteins has been demonstrated to be stepwise rather than continuous, with well-defined steps of ∼ 8 nm (*5*). These small steps require nanometer localization precision and fast and consistent sampling, which poses a significant challenge in RT-3D-SPT. Here unfunctionalized 100 nm AgNPs were tracked in Hela cells. The depolarized AgNP signal in cell media displayed a sufficient signal-to-background ratio for 3D-SMARTER tracking (Fig. S5), and this ratio was maintained over a large axial range (Fig. S6). In live HeLa cells, endocytosed AgNPs frequently exhibited linear trafficking behavior. Within these trajectories Fig. 3d-e), steps of 7.6 ± 2.5 nm were observed (Fig. 3f, step detection described in SI), comparable to prior studies (*39-42*).

**Figure 3.**
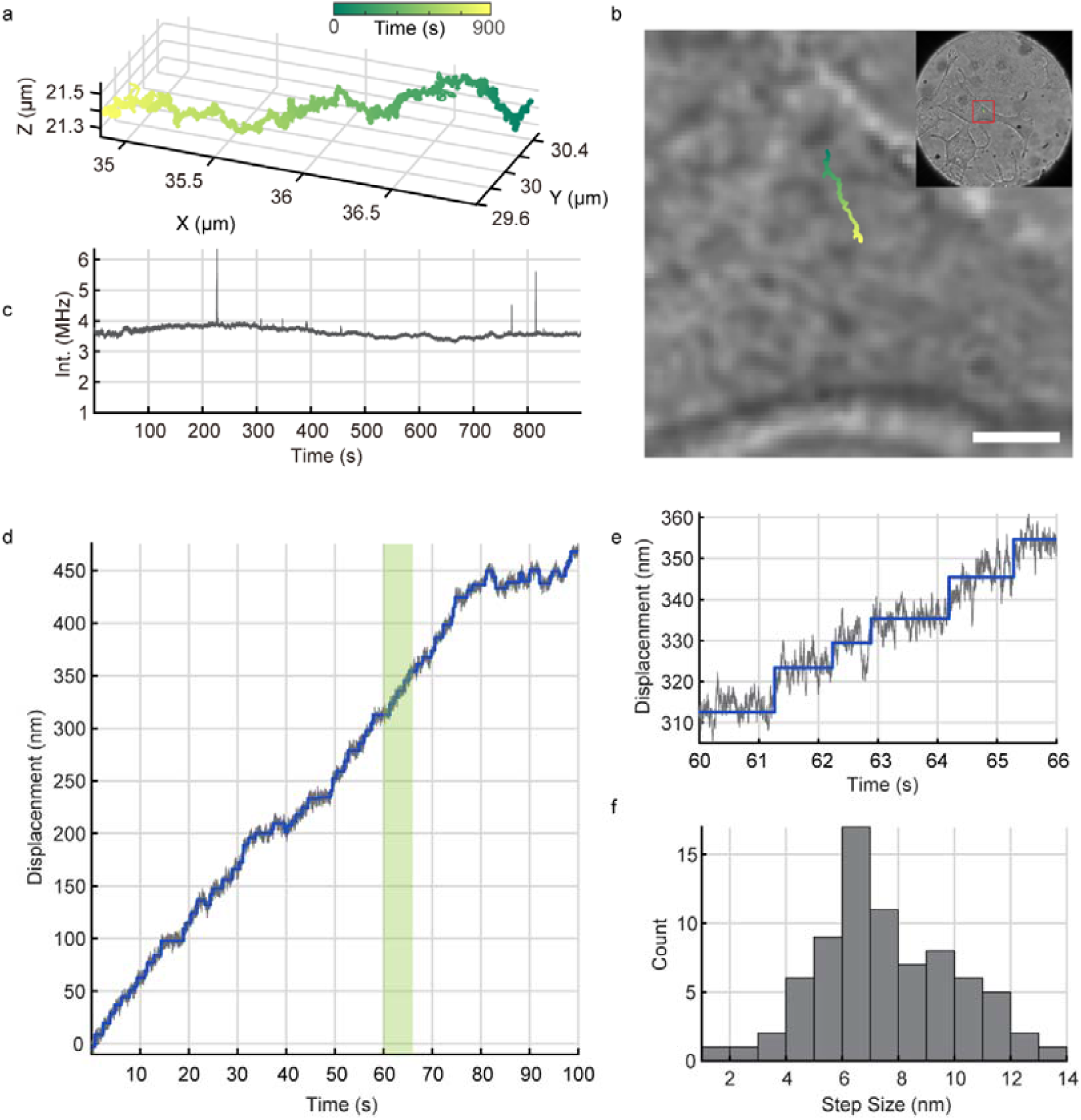
Characterization of motor protein motion in live cells by 3D-SMARTER. (a) 3D trajectory of a 100-nm AgNP in a live HeLa cell. (b) Bright-field image of cell overlapped with the trajectory in (a) obtained after the trajectory was collected by 3D-SMARTER. Scale bar: 2 um. Insert: larger view of cell. (c) Depolarized scattering intensity trace of the trajectory in (a). (d) Displacement in lateral motion along an axis parallel to the observed linear motion from 610-710s of the trajectory in (a). (e) A closer look of (d) with steps labeled by light green shaow. (f) Step size histogram. Mean step size = 7.6 ± 2.5 nm (n = 76).

### MHz localization in three dimensions with recursive Bayesian position estimates

In RT-3D-SPT, active-feedback is used to center the object within the observation volume in real time. If the feedback speed is faster than the process in question, then the current position of the piezoelectric stage should be an excellent approximation for the true particle position. For example, in results discussed above, stage positions sampled at 1 kHz were treated as the true particle positions. High-end piezo nanopositioners are bandwidth-limited to 1 kHz, which serves as a low-pass filter on the tracked particle position. This is not usually a barrier in fluorescent based RT-3D-SPT experiments, due to the fact that the fluorescence emission rate limits the sampling rate, rather than the mechanical response time of the stage. However, given the extremely high photon flux from the AgNP in these experiments, sampling far beyond this 1 kHz barrier is possible. Given the >3 MHz scattering rate, we were able to process photon arrivals into 1 µsec bins and use a recursive Bayesian analysis to untangle the true particle position from the relatively slow response of the piezoelectric stage (Fig. S7). The result is 3D particle tracking at 1 MHz sampling rate. The 1 μsec binned photons are used to localize the particle in three dimensions, treating XY and Z as linearly independent. Details of the calculation can be found in the SI. The reconstructed trajectory shows that the stage position is effectively a lowpass filter of the best estimate of the particle position (Fig. 4 a,d,g). The stage position lags particle motions estimated by the recursive Bayesian estimate, especially during fast motions (Fig. 4 b,e,f). Noticeably, the localization uncertainty of estimated X/Y position is comparable to that of Z position, despite of a larger PSF scale in Z and a doubled stage readout error, indicating similar information efficiency can be achieved among three dimensions.

**Figure 4.**
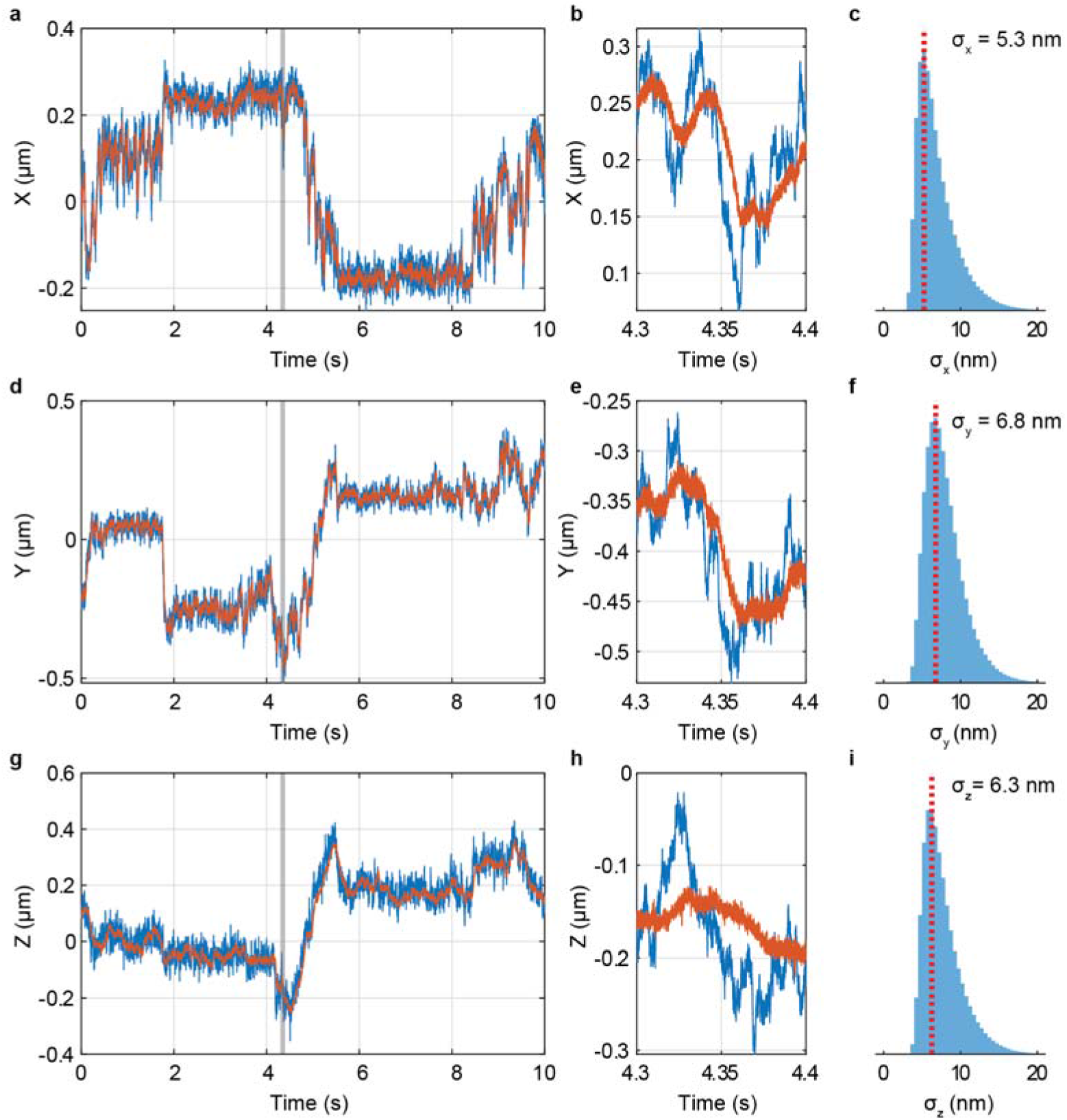
Recursive Bayesian position estimation enables single microsecond temporal resolution in 3D. (**a**,**d**,**g**) Stage positions (orange curve, sampled at 1 MHz) versus Bayesian estimates (blue curve, sampled at 1 MHz) of part of a freely diffusing AgNP trajectory. (**b**,**e**,**h**) Closer look of part of the amplitudes in (a,d,g). (**c**,**f**,**i**) Distribution of uncertainty from Bayesian estimates.

### AgNP tracking reveals interactions between AgNPs and local hot spots on HeLa cell filopodium

Filopodia are thin cell membrane protrusions that play an important role in cell adhesion, migration and endocytosis (*43, 44*). Filopodia also play an imporant role in viral infection, where viral “surfing” is frequently observed (*45-47*). Despite their extensive involvement in a variety of biological processes, the details of interactions between filopodia and cargos in live cells remain unclear due to the lack of methodologies that can visualize such interactions with nanometer and microsecond resolution. With 3D-SMARTER microscopy and AgNPs as bright and durable probes, it is possible to track cargo-filopodium interactions for a long observation time with high 3D spatiotemporal precision. Here we investigate confined particle diffusion on a cylindrical structure, including identification of diffusive “hot spots” and characterization of rotational movements. We found that cargo-filopodium interaction displays additional spatial complexity compared to previously discussed longitudinal diffusion driven by motor proteins. Here we utilized 3D-SMARTER to observe the interactions between HIV cell-penetrating TAT peptide (TATp) (*48, 49*) functionalized AgNP (TATp-AgNP) and HeLa cell filopodia. To functionalize the AgNP with TATp, streptavidin (SA) was first labeled with a thiol group (SH) via NHS-PEG-SH to give the protein an affinity for the AgNP surface. After mixing AgNPs with SA-SH, the resulting particles were incubated with TATp-biotin to functionalize the AgNPs with TATp. It was observed that, upon functionalization with TATp, AgNP had larger size (∼140 nm diameter functionalized compared to ∼100 nm diameter non functionalized) while the absorption spectrum remained unchanged (Fig. S8). TATp-AgNPs were added to live HeLa cells and then tracked by 3D-SMARTER. The positive charge of the TATp peptide gave the AgNPs an affinity for the cell membrane. It was observed that that higher concentration (56 μM) of TATp resulted in a higher incidence of intracellular trafficking events (19 out of 27) compared to lower TATp concentration (0.28 μM, 12 out of 55), as is shown in Fig. S10. This observation was consistent with previous studies that indicated nanoparticles with higher TATp valency showed increased tendency to undergo endocytosis (*50-52*). AgNPs labeled with arginine-9 (R9) peptides at high concentration comparable to TATp showed similar high percentage of internalization (18 out of 22). This further indicates that the fate of functionalized AgNP was dominated by surface charge.

Among bound particles, cylindrical diffusion on the surface of live HeLa cells was often observed. An example of such motion can be seen in Figure 5 for a single trajectory with a duration of over 20 minutes and average intensity of 2.4 MHz, carving out a cylindrical structure with a diameter of 346 ± 86 nm (Fig. S9a). Taking the diameter of the AgNP particle (∼100 nm) into account, the actual diameter of the tubular structure is ∼250 nm, consistent with the reported size of filopodial structures (*43, 53*). Part of the trajectory with 60 s duration length (900 s – 960 s in entire data set mentioned above) is shown in Fig. 5a. The cylindrical geometry is even more pronounced when the trajectory is projected onto a cylindrical coordinate system (*r*, radius; *θ* angle; *h*, height; Fig. 5b,c). The same trajectory segment observed from top of the cylinder in Fig. 5b is shown in Fig. 5h, and the 2D density of Fig. 5h is shown in Fig. 5i, which further illustrates the particle undergoing restricted diffusion on a cylindrical surface. A residence time analysis was performed to determine whether the particle was interacting with specific structures on the surface of the filopodium. The coordinates of the trajectory were first projected to the surface of a cylinder with a radius of the average *r* of the segment (Fig. 5d). The coordinates from 1 MHz Bayesian reconstruction were then converted to a 2D density map with 10 × 10 nm grids, as shows in Fig. 5e. The density map shows the particle covered most of the surface of the cylinder during diffusion while the residence time at different regions vary significantly with local “hot spots” (or areas of interest (AOIs)), were clearly observable as indicated with circles in Fig. 5e. Pixelated intensity in each AOI was approximated as a 2D Gaussian distribution defined as:

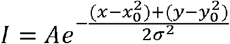

Here *I* is the dwell time of the particle at a given pixel, *A* is a prefactor, *x*_0_ and *y*_0_ are the center coordinate of the Gaussian spot and is the uncertainty of the spot. Fig. 5f-g show the brightest AOI in Fig. 5e and its 2D Gaussian fitting image. The border of a given AOI is defined as a circle within (95% confidence interval) of the center of the AOI in the 2D density map. The 98 manually classified “hot spots” had an average size of 55 ± 14 nm and were visited a total of 140 times, with 29 “hot spots” being visited multiple (≥ 2) times. The repeated visits in AOIs indicates the dwelling events could originate from potential structures interacting with the AgNP. The residence time distribution fits well with a 2-factor exponential decay with half-lives of 0.28 and 5.96 s (Fig. S11). Notably, the events with shorter duration were likely from purely diffusive, non-binding events.

**Figure 5.**
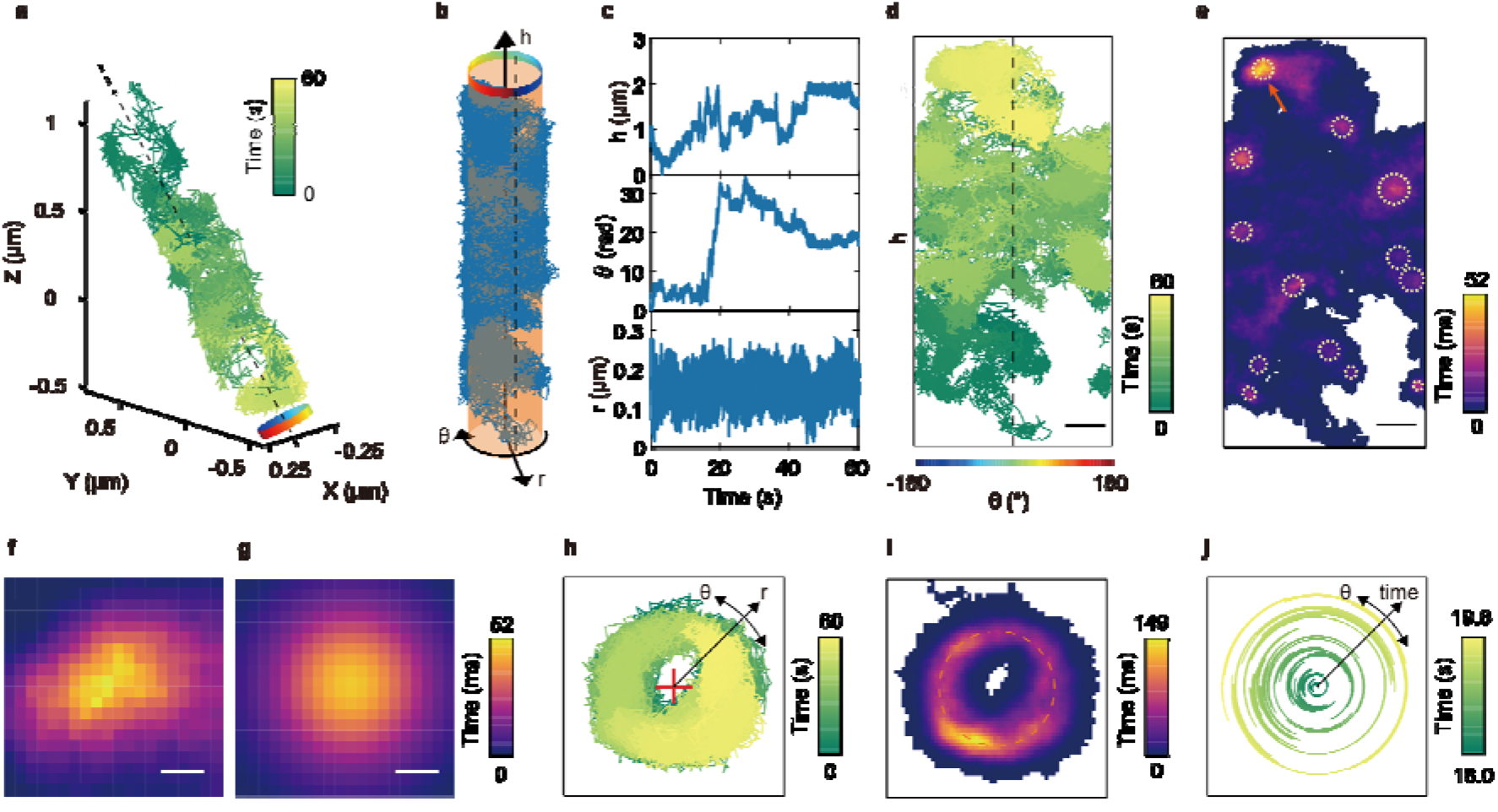
Mapping the nanoscale structure of filopodium with microsecond resolution by 3D-SMARTER. (**a**) 60-second of trajectory (sampled at 500 Hz) showing restricted diffusion on a cylindrical structure. (**b**) Demonstration of cylindrical coordinate system overlapped with the trajectory in (a). (**c**), and versus time from the trajectory in (a). Data sampled at 1 kHz. was expanded beyond ±180° for smooth visualization. (**d**) 2D trajectory of the trajectory in (a) projected onto the cylinder shown in (b) with a radius of 157 nm (average of shown in (c)). Dashed line is consistent with the dashed line in (b) and color bar on the bottom showing angle was consistent with the curved color bars in (a) and (b). Data sampled at 1 kHz. (**e**) Density map of residence time of 2D trajectory sampled at 1MHz. Dashed circles show AOIs. (**f**) Residence time distribution of AOI labeled with orange arrow in (e). (**g**) 2D Gaussian fit with a of 50 nm of data shown in (f). Scale bars in (f) and (g): 50 nm. (**h**)Trajectory observed from top of *h* axis in (b). Data sampled at 1kHz. (**i**) Density map of residence time of 2D trajectory shown in (h) sampled at 1MHz. (**j**) Angle versus time of part of the trajectory in (a). Growing radius of the helix plot indicates time progress.

We also investigated the speed and direction of rotational (motion in direction) diffusion on filopodium. Fig. 5j shows the angular position as a function of time, where the particle underwent fast and clockwise rotation over 3.8 s. After labeling and analyzing fast rotation events in 13 intervals, it was found that there was no significant difference in the speed and directionality of rotations, with 52 clockwise events of 13.5 ± 6.4 rads/s and 60 counter-clockwise events of 15.3 ± 7.5 rads/s (Fig. S9b-d). The complete trajectory shown in Fig. 5 overlapped with the cell was shown in Fig. S12. This indicates that the cylindrical motion was random, rather than biased in any particular direction.

### Surface force mapping shows local potential wells overlapped with high residence time AOIs

With sufficient spatial and temporal resolution, it is possible to extract more information than just the local diffusion coefficient from trajectories. A moving particle on a complex surface will experience areas of different viscosity, physical barriers, and local potentials. Measuring these local potentials requires localization and temporal precision on the scale of the potentials involved. The spatiotemporal precision enabled by 3D-SMARTER here allows mapping of forces on complex 3D surfaces with nanoscale resolution. The process applied here is derived from a Bayesian algorithm developed by Masson and co-workers (*54*). Briefly, to measure both the isotropic diffusion and local drift velocity simultaneously, the data are gridded, and the following likelihood is applied to each gridded position:

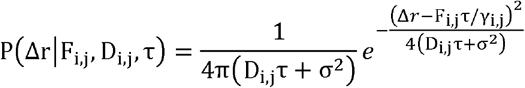

The equation gives the probability of observing a displacement (Δ*r*) given a particular grid position (*i, j*), delay time (τ), local diffusion coefficient (*D*_*i,j*_), local force (*F*_*i,j*_), local drag coefficient (γ_*i,j*_), and measurement noise (σ). By multiplying the probability for each observed displacement, a maximum likelihood estimate (MLE) can be extracted for both the local diffusion coefficient *D*_*i,j*_ and local vector forces *F*_*i,j*_, which can be used to extract the scalar potential. The results of this analysis on a 60 s segment of filopodial surface diffusion is shown in Fig. 6. There is striking difference between the map of the MLE estimated diffusion coefficients (Fig. 6a, c) and the density of spots at each gridded position (Fig. 6b, d). There seems to be no correlation between the diffusion coefficient and the amount of time a particle spends in a particular hotspot. In contrast, the MLE estimated force-vectors show that particles are constrained to nanoscale domains due to directional motion which forces the particle back into small areas. This indicates that these hot spots are not simply pockets of altered viscosity but are actual potential wells which trap the particle in a particular domain. Surface integration of the force vectors allows a full reconstruction of the nanoscale potentials on the surface of the filipodium, as shown in Fig. 6e.

**Figure 6.**
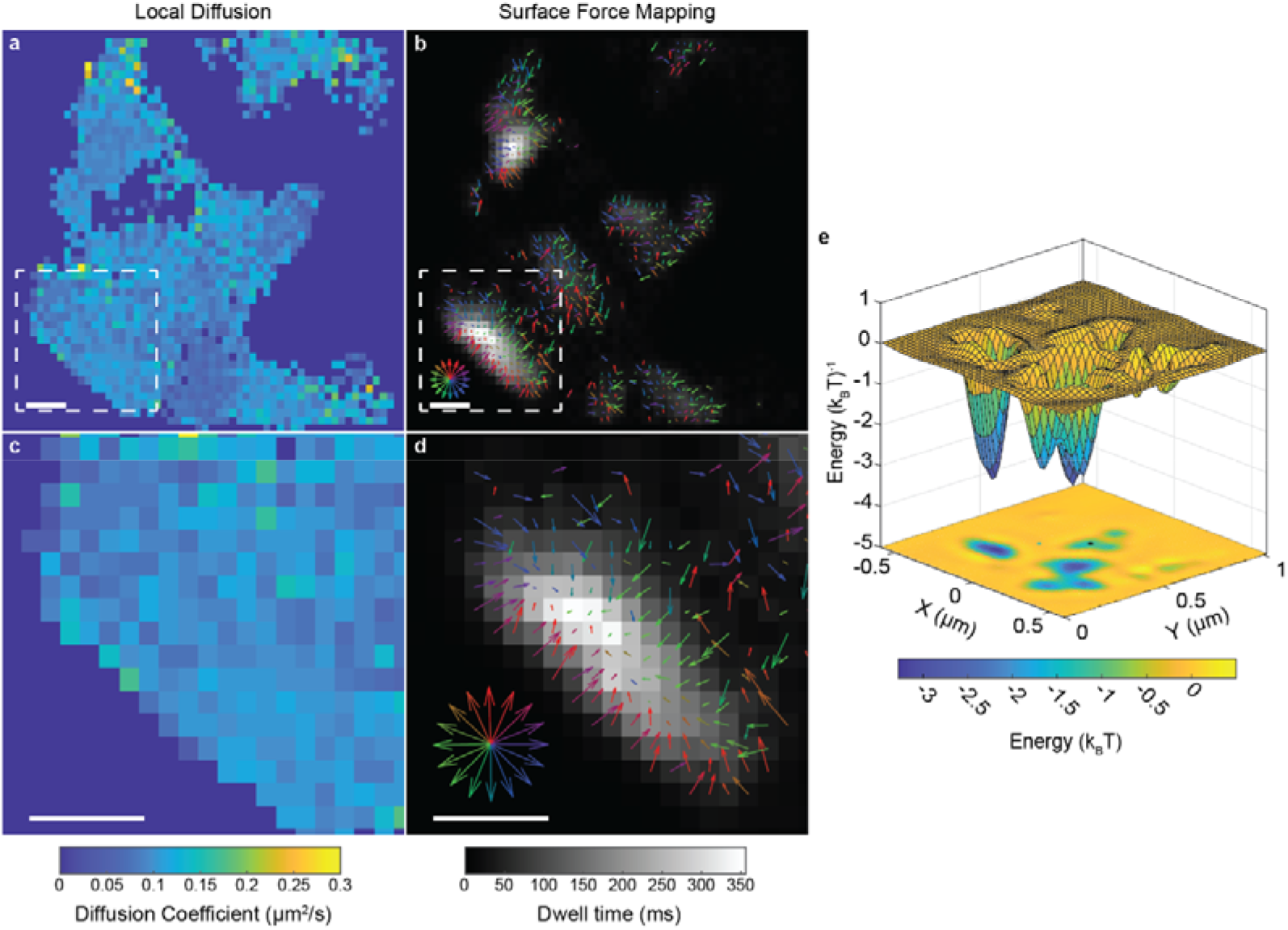
Extracting force-vectors and nanoscale potential wells on 3D surfaces. (**a**) Map of the MLE diffusion coefficient on the surface of a filopodium, carved out by the motion of a single AgNP, localized at 1 µsec intervals. (**b**) Zoom in of square area shown in top panel. (**c**) MLE extracted force-vectors (arrows) overlaid with the dwell time at each grid position (grayscale). (**d**) Zoom in of square area shown in top panel. (**e**) Potential surface on the surface of a nanoscale filopodium. Scale bars: 100 nm. Scale arrows: 0.1 pN.

## Summary

In this study, we introduce 3D-SMARTER, a real-time tracking method utilizing depolarized detection of scattering AgNPs. The combination of AgNPs and rapid feedback tracking of 3D-SMART provides a large sampling depth of over 50 μm, spatial precision down to 2 - 4 nm in XYZ, and temporal sampling down to 1 μsec. Critically, the microsecond temporal resolution and nanometer localization precision can be extracted from a single trajectory with a duration time over tens of minutes in a live cell. The combined spatiotemporal precision enabled mapping nanoscale hotspots and potential wells on the surface of cylindrical filopodia. These studies show the power of high spatial and temporal precision of 3D-SMARTER. When temporal sampling is poor, it is only possible to extract changes in diffusion coefficient, and even then these changes are limited by the spatial resolution of each spot. Here, 3D-SMARTER goes far beyond simple changes in diffusion coefficient, or deviations in the linearity of a mean-square displacement analysis. Drift velocities, their associated force vectors, and even potentials can be extracted with nanoscale precision. 3D-SMARTER has the potential to provide potential energy surfaces on a wide range of biological structures, yielding an ever-clearer picture of forces at the nanoscale in biology. In future studies, dark-field illumination can be employed to achieve an even higher signal-to-background ratio, leading to 3D-SMARTER for smaller particles and with better spatial precision. Anisotropic scattering probes, such as nanorods, can also be explored in 3D-SMARTER to study the rotational potentials on complex biological surfaces (*26, 55-57*), adding yet another dimension to this powerful method.

