## Supplementary Methods and Data for "Mapping nanoscale forces and potentials in live cells with microsecond 3D single-particle tracking"

Hou et. al.

#### Functionalization of AgNPs with TATp and R9 peptides

Biotin-TAT peptides (AS-61209) and R9 peptides (AS-64078) were purchased from AnaSpec, Inc. First, streptavidin (SA, Z7041, Promega Corp.) was labeled by NHS-PEG-Thiol (PG2-NSTH-3k, Nanocs Inc.) using following steps: (1) mix SA (36 μM) with NHS-PEG-Thiol (200 μM) in PBS solution; (2) incubate the solution in 25 ˚C for 2 hours; (3) purify the labeled SA with NAP-5 G25 desalting column (17085301, GE Healthcare illustra) following the product manual. The SA-SH solution (9 μM) was then mixed with 100-nm AgNPs (165 pM, AGCB100-1M, NanoComposix) and incubated at 25 ˚C for 2h. The SA-AgNP solution was then mixed with biotin-TAT peptides (2.8 μM for low labelling density and 56 μM for high labelling density) or biotin-R9 peptides (56 μM) and incubated at 25 ˚C for 2h. The solution was then purified by centrifugation (1600g, 20 min).

#### Tracking TATp functionalized AgNPs in live HeLa cells

HeLa cells were cultured on coverslip for 12h before the tracking experiment. Then cell culture medium was washed 3 times and replaced with live cell imaging solution (A14291DJ, Molecular Probes). Then ~3pM AgNPs were added into the cell solution for real-time tracking. Laser power was 0.64 μW.

#### Measurement of signal-to-background ratio with scattering images

Images used for signal-to-background measurement were taken from raster scanning with piezo stage with pixel dwell time of 10 ms. A scattering image was first taken before adding the TATp functionalized AgNPs. Then a second image of the same area was taken after the 3 pM AgNP solution was added.

#### Tracking the intracellular trafficking of unfunctionalized AgNPs

The cells were cultured on coverslip for 12h before the tracking experiment. The cell culture medium was then washed with live cell imaging solution (A14291DJ, Molecular Probes) three times and resuspended in the imaging solution for live-cell imaging. Then ~3 pM nonfunctionalized AgNPs were added to cell solution for tracking. The laser power for tracking was 0.64 μW.

#### Step identification in intracellular trafficking

The entire trajectory of internalized AgNPs was first split into 3-second segments. Each segment was fit to line to determine the overall displacement vector. The datapoints for each segment were then projected to the corresponded vectors and produced the scalar displacements. Then the scalar displacements for all segments were combined. Steps were identified and labelled using algorithm developed by Kerssemakers and coworkers (Nature 2006, 442, 709–712).

#### Colocalization of TATp functionalized AgNPs and actin

Actin was stained with SiR-Actin Kit (CY-SC001, Cytoskeleton, Inc). SiR-Actin was diluted into 1 μM in cell culture medium and added to cell solution. Cells were then incubated for 1 hour in cell incubator. After staining, cells were washed 3 times with cell imaging buffer and then TATp functionalized AgNPs (~3 pM) were added into cells for live cell tracking. 640-nm laser was used to obtain the fluorescent image of actin. A dichroic filter (T610lpxr, Chroma) was used to separate actin fluorescence (excited by 640 nm laser) and AgNP scattering signal (488 nm laser). A bandpass filter (ET706/95m, Chroma) separates the fluorescence emission from the excitation lasers. 3D fluorescent images of actin were obtained through piezo stage guided raster scan of cell sample immediately after single particle tracking was complete.

**Recursive Bayesian Localization for MHz sampling of 3D SMARTER trajectories**

The first step for recursive Bayesian localization of AgNPs was to bin collected photon arrival times, initially collected with 12.5 nsec resolution, into 1 usec bins. Each photon arrival was tagged with the current piezoelectric stage coordinates and laser scan position. They XY and Z data were treated independently. For XY, the expected observed photon emission rate at a given laser position is:

$$\gamma(x,y) = s e^{\frac{-(x-x_{laser})^2}{2\sigma^2}} e^{\frac{-(y-y_{laser})^2}{2\sigma^2}} + b \tag{1}$$

Here, $\gamma(x,y)$ is the expected emission rate for a particle of brightness $s$ and background $b$ located at position $(x,y)$, illuminated by a laser at position $(x_{laser}, y_{laser})$ with beam width $\sigma$. The probability of observing a number of photons $n$ is determined by a Poisson distribution:

$$P(n|x,y) = \frac{(\gamma(x,y)\tau)^n e^{-\gamma(x,y)\tau}}{n!} \tag{2}$$

The maximum of this function yields the maximum likelihood estimate (MLE) of the particle position. In recursive Bayesian inference, the posterior likelihood distribution is

$$P(x,y|n) = \frac{P(n|x,y)P(x,y)}{P(n)} \tag{3}$$

P $P(x,y|n)$ is the posterior distribution, $P(n|x,y)$ is the likelihood of observing the current number of photons $n$, $P(x,y)$ is the prior distribution, and $P(n)$ is the probability of observing $n$ photons. In log form:

$$log\ P(x,y|n)\ =\ log\ P(n|x,y) + logP(x,y) - log(P(n)) \quad (4)$$

In practice, the final term serves to normalize the posterior distribution. The unnormalized log-posterior distributions is:

$$log\ P(x,y|n)\ =\ log\ P(n|x,y) + logP(x,y) \quad (5)$$

Or

$$log\ P(x,y|n)\ = logP(x,y) + nlog(\gamma(x,y)\tau) - \gamma(x,y)\tau - log(n!) \quad (6)$$

So the log-posterior is the sum of the log-prior and the current log likelihood. For a non-moving particle, this can be used to localize a particle by recursively updating the prior distribution with the likelihood of each arriving photon, starting with an initialized (uniform) prior and normalizing the posterior distribution at each step.

An additional step is needed for a particle that moves. If the particle is diffusing, one cannot simply assign the posterior distribution from the previous step to become the prior for the current observation. To account for this, the normalized posterior is convolved with a a diffusion kernel:

$$\iint P(x,y)\, e^{-\frac{(x-x')^2+(y-y')^2}{4D\tau}} dx'dy' \quad (7)$$

$D$ is the expected diffusion coefficient.

1. Photon arrivals, along with concurrent laser and stage positions, are binned into one µsec intervals.

2. An initialized and uniform prior distribution is generated.

3. Starting with the first bin, Equation 2 is used to generate the current likelihood.

4. The posterior distribution is calculated using Equation 6 using the prior (step 2) and current likelihood (step 3).

5. The posterior is convolved with the diffusion kernel via Equation 7.

6. The posterior becomes the new prior and the process goes back to step 3 and proceeds through the entire dataset.

**Supplementary Figure 1: 3D-SMARTER setup**

The setup of 3D-SMARTER is similar to previously reported 3D-SMART with a linear polarizer added in the detection path. Briefly, A 488-nm frequency-doubled solid-state laser (FCD488-30, JDSU) is used for the excitation of AgNPs and a 640-nm diode laser (OBIS 640LX, Coherent) is used for the excitation of actin fluorescent imaging. The polarized of the laser beam is cleaned by a polarizer and tuned by a half-wave plate. The laser is then deflected by a pair of electro-optic deflectors (EOD; M310A, ConOptics) to generate a designated information-efficient excitation pattern in the XY-plane. Then a tunable acoustic gradient lens (TAG Lens 2.5, TAG optics) is used to perform varifocal excitation in Z-axis. To achieve real-time active feedback 3D tracking, a 3D piezoelectric stage system (XY: Nano-PDQ275HS, Z: Nano-OP65HS, Mad City Labs) is used to move the sample to keep the particle being tracked in the center of the laser sampling volume. The excitation light and fluorescence light are splitted with a dichroic mirror (ZT405/488/561/640rpc, Chroma). It should be noted here that this dichroic mirror will also filter out most of the scattering signal. The fluorescent signal and scattering signal are split by a dichroic mirror (T510lpxru, Chroma). A field-programmable gate array (FPGA, NI-7852r National Instruments) is used to control the excitation pattern generated via EODs and TAG lens, measure the particle position in real time, apply active position feedback with the piezoelectric stage, and record particle position information. In the detection path of scattering tracking, a polarizer and a bandpass filter (ZET488/10x, Chroma) are used to extract the depolarized scattering signal. In the detection path of fluorescent imaging, a bandpass filter (ET706/95m, Chroma) is used to separate the fluorescent emission from the excitation laser. Two avalanche photodiodes (APDs) are used to measure the scattering signal and fluorescence signal, respectively.

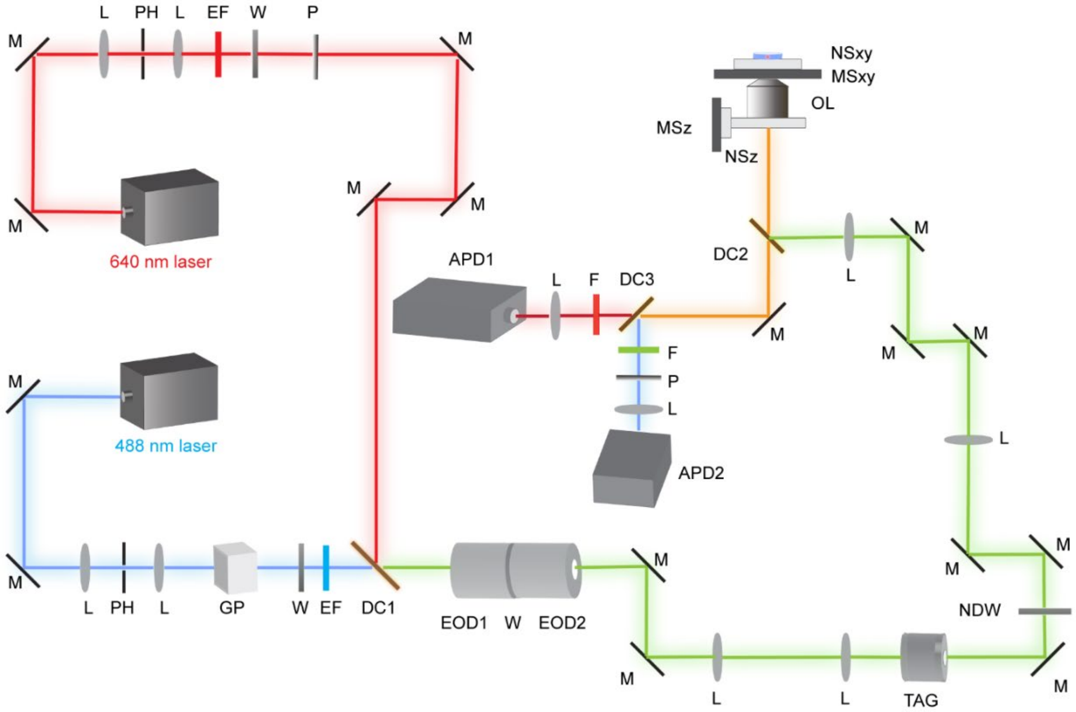

**Figure S1.** Schematic of 3D-SMARTER setup. L: lens; M: mirror; PH: pinhole; GP: Glan-Thompson polarizer; P: polarizer; W: half waveplate; EF: excitation filter; DC: dichroic mirror; EOD: electro-optic deflector; TAG: TAG lens; NDW: Neutral density filter wheel; OL: objective lens; MSxy: XY microstage; MSz: Z microstage; NSxy: XY nanopositioner stage; NSz: Z nanopositioner stage; F: fluorescence filter.

### Supplementary Figure 2: Information-efficient excitation pattern

In the previously reported 3D-SMART setup, the laser spot was guided to continuously sample a 5*5 grid in the XY plane through Knight's Tour pattern (Fig. S2a). In this study, a 4-pixel custom pattern named '4-Corners' pattern (Fig. S2b) was used to sample the XY plane as previous study showed that Fisher information (FI), which determines the theoretical lower bound of precision from a measurement, is maximized at a certain distance from the center of the emitter in single dimension. Specifically, when the intensity of a diffraction-limited emitter at x is approximated as Gaussian with center of μ and uncertainty of σ in arbitrary dimension:

$$f(x|\mu,\sigma) = \frac{1}{\sqrt{2\pi\sigma^2}} e^{\frac{-(x-\mu)^2}{2\sigma^2}}$$

The FI of the emitter can be derived as:

$$\mathrm{J}(\mu) = \int f(x|\mu) \frac{(x-\mu)^2}{\sigma^4} dx$$

This gives maximum FI at $\pm\sqrt{2}\,\sigma$ from the center of the emitter. It is also noticeable that there is no FI at the center of the particle (Fig. S2c). In the XY-plane, the 4-Corners pattern overlaps with the high FI area (Fig. S2d). In the Z-axis, the laser is guided to scan following a sine wave with frequency of ~70kHz (Fig. S2e) as previously reported, as the unmodulated sine wave pattern display information efficiency due to the intrinsic edged distribution of probability density function of the sine wave (Fig. S2f).

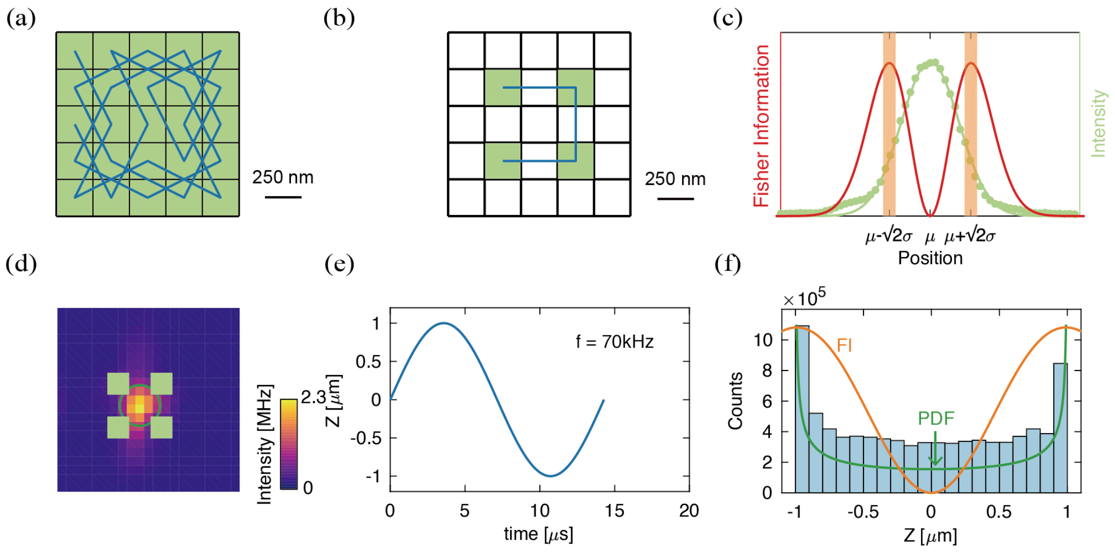

**Figure S2.** (a) XY scanning pattern of previously reported 3D-SMART microscope. (b) XY scanning pattern of 3D-SMARTER. (c) Fisher information distribution (red line) of Gaussian approximate (green line) of experimentally obtained intensity along X-axis from a fluorescent particle (green dots). (d) High FI positions (green circle) and proposed sampling positions of the 4-Corners pattern. (e) Z position versus time for TAG lens. (f) Photon arrival distribution along Z-axis from the trajectory of a freely diffusing 100-nm AgNP in water.

**Supplementary Figure 3: Measurement of Signal-to-background value at different polarizer angles**

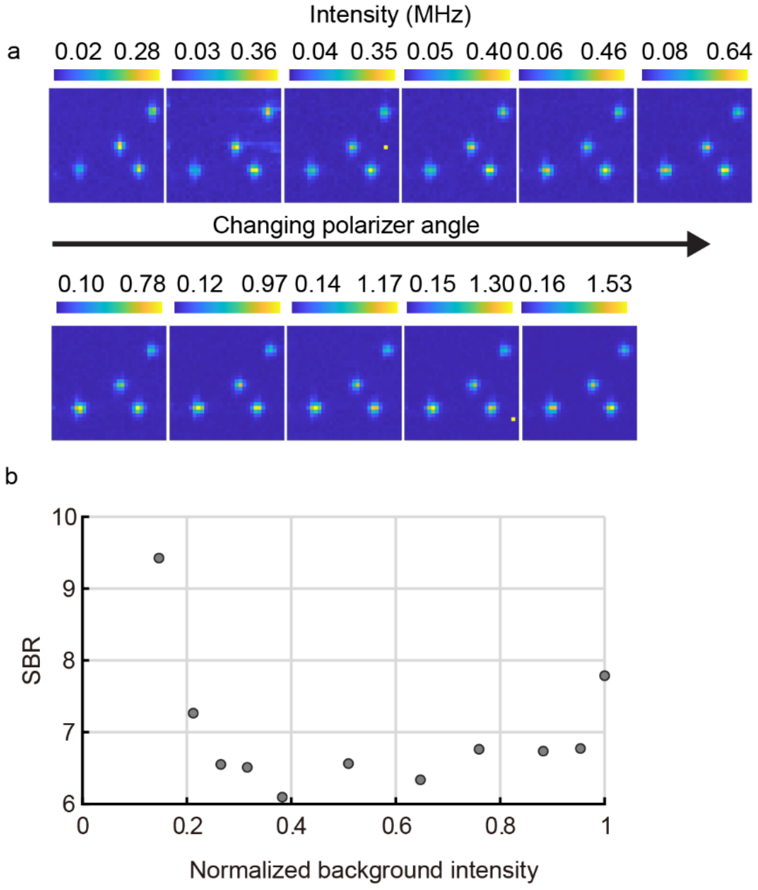

**Figure S3.** (a) AgNP scanning images obtained from continuously changing polarizer angles. Scanning size: 31 × 31 pixels. Pixel size: 200 nm. Laser power: 488 nm, 6.4nW. (b)Signal to background ratio (SBR) as a function of polarizer angle. Signal is calculated as the average intensity of the four particles and the background is calculated as the average intensity of an area of 6 by 6 pixels with no particle present.

**Supplementary Figure 4: Tracking of freely diffusing AgNP with 3D-SMARTER**

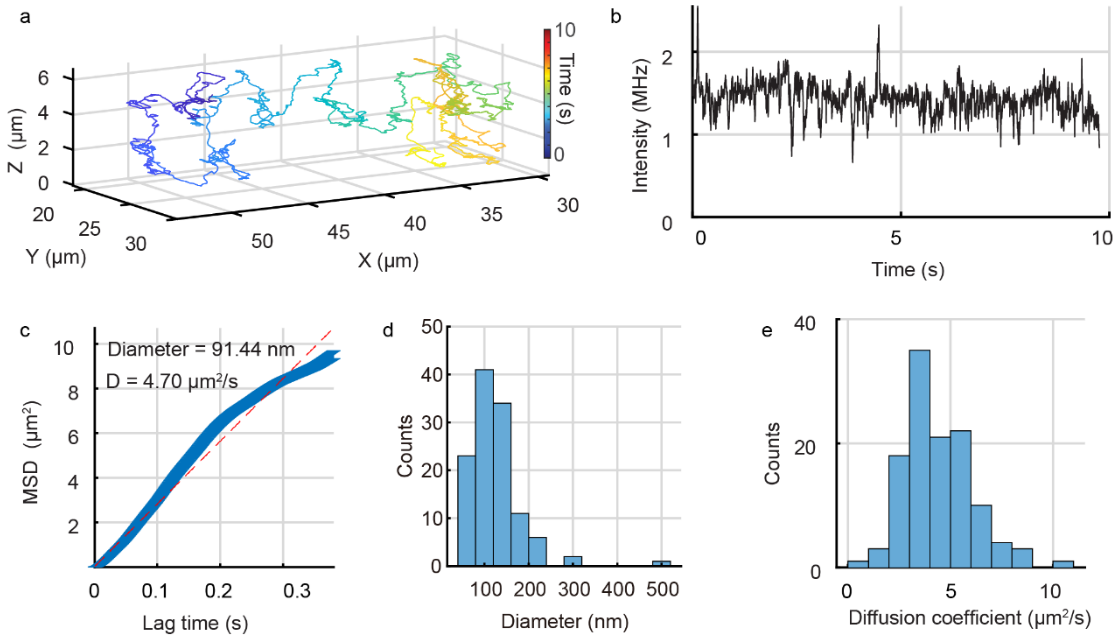

**Figure S4.** (a) 3D trajectory of a 100-nm AgNP freely diffusing in water. (b) Depolarized scattering intensity trace of trajectory in (a). (c) Mean square displacement versus lag time in trajectory in (a). (d) Distribution of estimated particle diameter of AgNPs tracked in water. Bin width= 40 nm.. (e) Distribution of diffusion coefficient of tracked AgNPs. Bin width = 1 $\mu m^2/s$. D = 4.43 ± 1.71 $\mu m^2/s$; Diameter = 124.63 ± 59.07 nm; n= 118; 100-nm PEGylated AgNPs (KJW1912, NanoComposix) were diluted in water (2 pM).

**Supplementary Figure 5: Images of live HeLa cells before and after introduction of TATp labeled AgNPs**

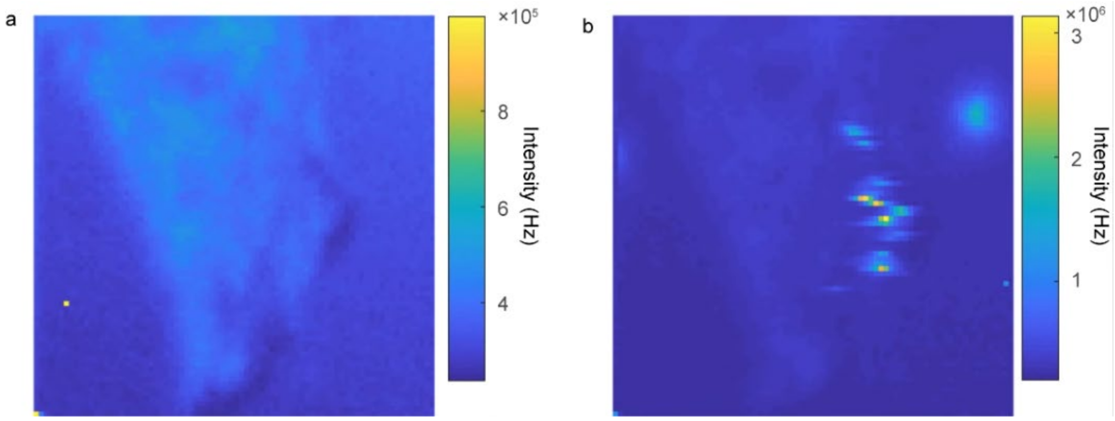

**Figure S5.** Scattering image before (a) and after (b) TATp labelled AgNPs added. Pixel size: 500 nm. Imaging range: 40 × 40 μm.

### Supplementary Figure 6: Signal-to-background measurement of AgNP tracking in HeLa cells

Though signal-to-background contrast was optimized in scanning image of immobilized AgNP in Fig. S3, it is important to measure signal-to-background value from tracking TATp functionalized AgNP signal in live Hela cell as cell medium is a more complex system with additional source of background. To achieve this, the cell medium loaded with TATp functionalized AgNP was first tracked with 3D-SMARTER. Once a particle was located and successfully tracked, tracking program was disabled and stage was guided to perform a raster scan with scattering signal consistently recorded. When the stage was off the particle, intensity profile was treated as cell background signal. It is noticeable that there are bursts in Fig. S6(b) where intensity was not comparable to tracking signal. This was due to the laser spot was partially overlapped with the particle, resulting in lower intensity compared to active tracking, when laser spot was highly overlapped with the particle.

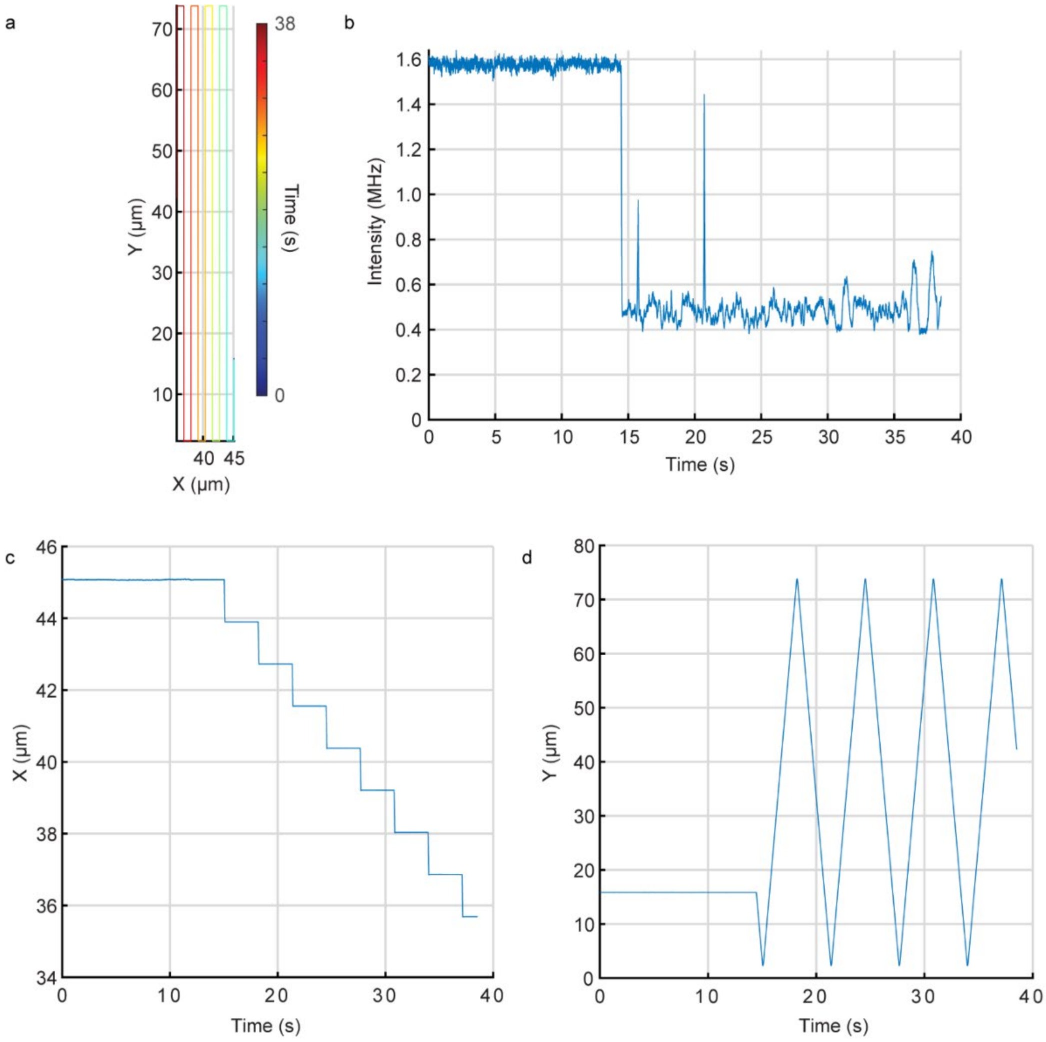

**Figure S6.** (a) Raster scanning pattern enabled by piezo stage. (b) Scattering signal during tracking (from 0 s to 14.4 s) and raster scanning (from 14.3 s to end). (c, d) X and Y position during tracking and raster scanning.

**Supplementary Figure 7: Simulated post-processing data of a Kalman tracked particle using photon arrival information**

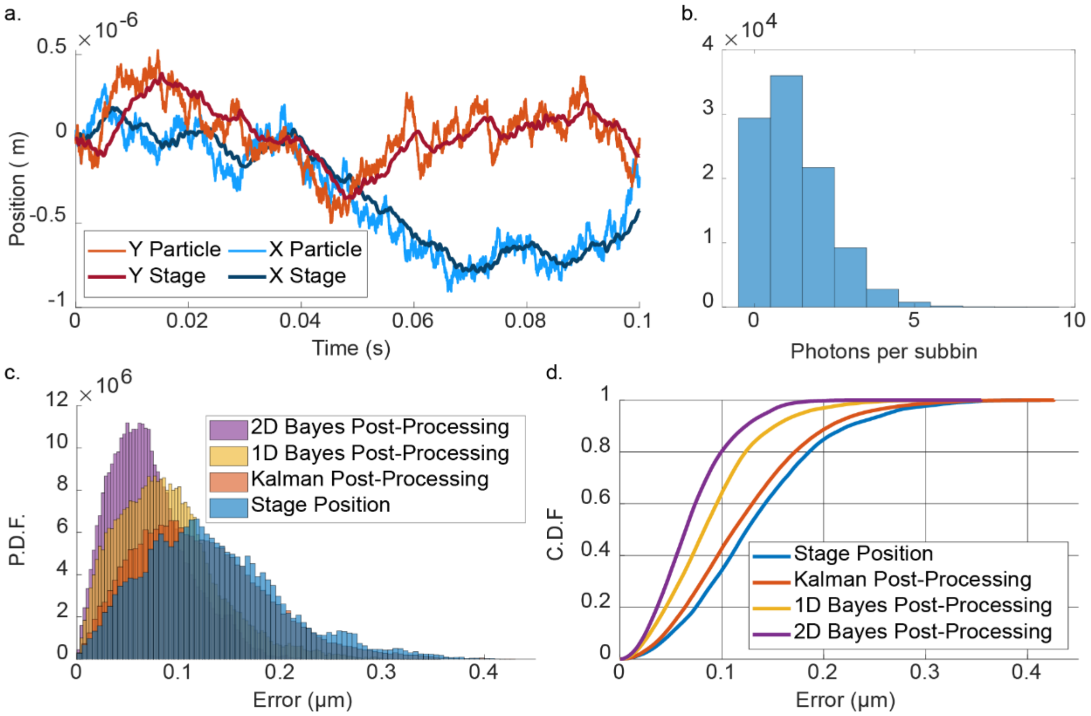

**Figure S7. Simulated post-processing data of a Kalman tracked particle using photon arrival information** a. Simulated particle positions and corresponding stage positions for a 2D trajectory of AgNP like particle tracked using experimental conditions. The stage lags the particle's true position. b. Photon arrival counts per 1 µs bins, used for post-processing. c. probability density normalized histograms of the error of 3 post processing algorithms as well as the original stage position (assumed to be the particle position in analysis without post-processing). Means are 0.133, 0.121, 0.088, and 0.070 µm for stage position, Kalman, 1D Bayes, and 2D Bayes respectively. b. Empirical cumulative density function representation of the same 4 errors in which it is easier to see that the 2D Bayesian post-processing in addition to improving the temporal resolution of the trajectories also approximately halves the error in actual particle position. Medians are 0.125, 0.112, 0.083, and 0.065 µm for stage position, Kalman, 1D Bayes, and 2D Bayes respectively.

### Supplementary Figure 8: Characterization of Functionalized AgNPs

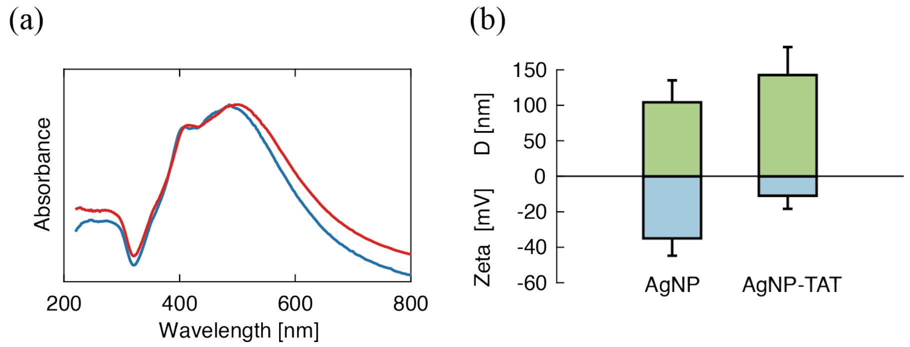

**Figure S8.** (a) UV-vis absorption spectrum of non-functionalized (blue curve) and TATp-functionalized AgNPs (red curve). Absobanse readout was normalized for comparison. Functionalized AgNPs were incubated with low density of TATp (2.8 μM). (b) Diameter (D) measurement from dynamic light scattering and zeta potential (Zeta) measurement of non-functionalized AgNPs and TATp-functionalized AgNPs (AgNP-TAT). Diameter was measured to be 104 ± 31 nm for AgNP and 141 ± 39 nm for AgNP-TAT. Zeta potential was -35.1 ± 9.7 mV for AgNP and -11.0 ± 7.4 mV for AgNP-TAT. Each value was from 4 measurements.

### Supplementary Figure 9: Analysis of rotation direction

Radius coordinates from cylindrical systems in 13 individual intervals showed the particle held a r position of 173 ± 43 nm from center axis (Fig. S9a). To investigate if rotation was biased along a certain direction, fast rotation events were manually labelled and all segments were observed 'top-down', as shown by arrows in Fig. S9b, to determine if individual rotation events were clockwise (CW) or counter-clockwise (CCW). An analysis of 112 labeled fast rotation events from 13 intervals showed that there are 52 CW events and 60 CCW events, indicating that such rotation is likely to be random and not biased along a certain direction. Rotation velocity of fast rotation events was found to be 12.0 ± 8.3 rads/s for CW events and 12.6 ± 5.4 rads/s for CCW events, which is also comparable along different direction. It was noticeable that all the fast rotation events from different time intervals are analyzed collectively with the assumpsion that the particle was interacting with the same structure during the entire trajectory as the center axis of the cylinders show a slowly wavering pattern (Fig. S9c). An example of manually labeled fast rotation events is shown in Fig. S9d.

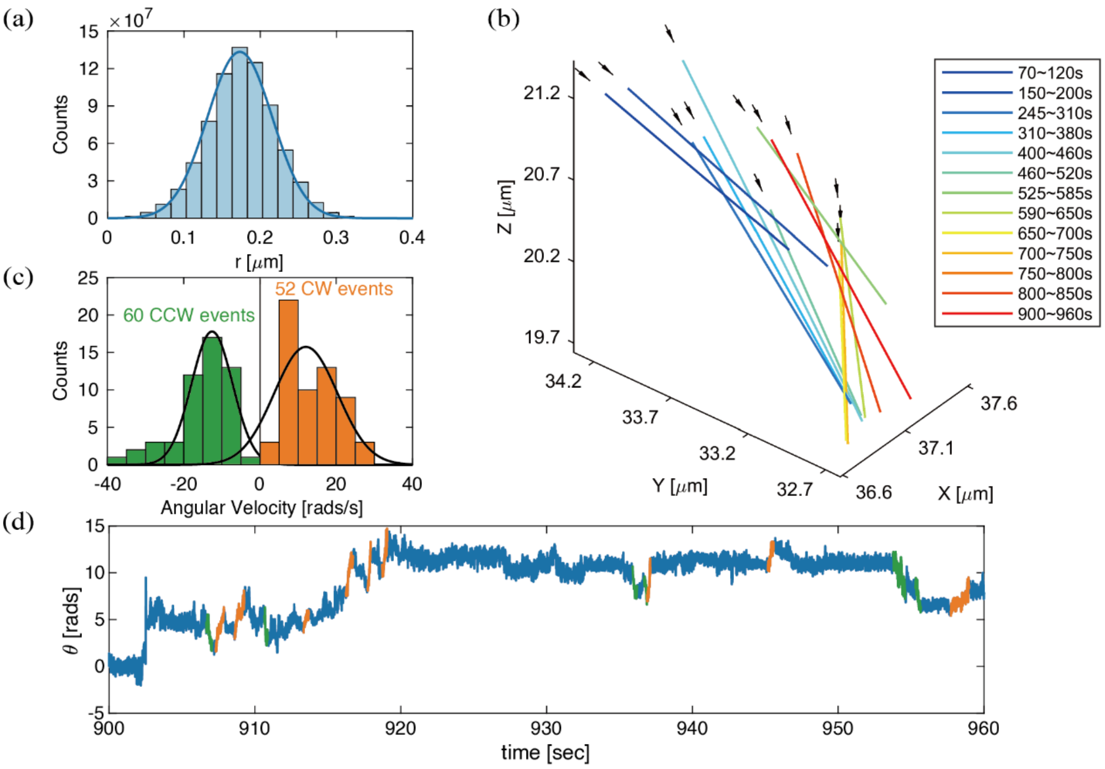

**Figure S9.** (a) Distribution and Gaussian fit of radius coordinates from all data points in 13 intervals. (b) Center axis of cylinders in 13 different intervals and direction of observation was labeled by arrows for individual axis. (c) Distribution and Gaussian fit of rotation velocity of 112 fast rotation events. Gaussian fit was performed separately on events in different rotations (CW or CCW).(d) labeled fast rotation events (orange lines and green lines stand for CW and CCW events, respectively) for data in 900-960s interval. Intervals with r<50 nm were not analyzed as those could result in abrupt change in angle upon projected to cylinder surface.

**Supplementary Figure 10: Different behavior of interactions observed from different types of surface functionalization of AgNPs**

It was noticed that functionalized AgNPs display different behaviors interacting with cells with different surface functionalization and at different functionalization density. By sorting trajectories from TATp-functionalized AgNPs interacting with cells, we found that higher concentration of TATp resulted in higher ratio of cellular trafficking trajectories (19 out of 27 display trafficking events) compared to lower TATp concentration (12 out of 55 display trafficking events), as is shown in Fig. S10. This observation was consistent with previous studies that indicated nanoparticles with higher TAT valency showed increased tendency to undergo endocytocis (Suzuki-2013). AgNPs labeled with arginine-9 (R9) peptides at high concentration comparable to TAT showed similar high percentage of internalization (18 out of 22). This further indicates that the fate of functionalized AgNP was dominated by surface charge.

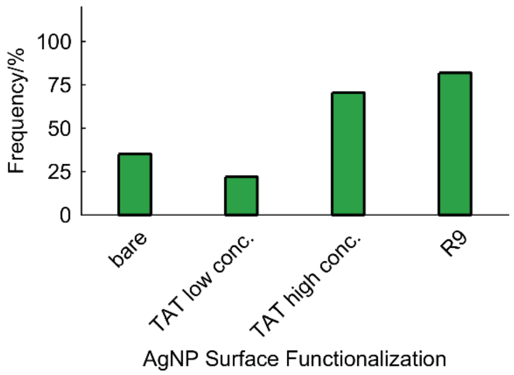

**Figure S10.** Percentage of trajectories associated with endocytosis behaviors observed from AgNPs with different surface functionalization interacting with HeLa cells.

### Supplementary Figure 11: Residence time analysis

Each manually labeled AOI was first evaluated through a morphology test which evaluates the distribution of visited positions along the $r$ axis. Number of visited positions (1 MHz Bayesian reconstructed data) inside a sphere with a radius of $\sigma$ the of each AOI along the $r$ axis was evaluated and the $r$ position where maximum data points were found along with $\theta$ and $h$ that were already available with AOI Gaussian fitting, were defined as center of an AOI. Only AOIs with a center coordinate $r$ inside the range of 87~259 nm ($\mu \pm 2\sigma$ of global $r$ distribution shown in Fig. S9a) were used for such analysis. Also those with number of spots found at $r = 0$ larger than 50% of points found at center $r$ were not considered in further analysis as such distribution could imply an abnormally high portion of spots found at center of the cylinder and thus made it unlikely for the particle to diffuse on a hollowed cylindrical structure. Of 132 AOIs manually labelled, 88 passed the morphology criteria described above, indicating most AOIs correspond to particle found frequently circulating at the surface of a cylindrical structure, which was consistent to the model of cargo interacting with unknown structures brought out in the article. To evaluate the binding dynamics of AgNP and filopodium, we divided 13 time intervals of 50~60s duration into 100-ms intervals and define that when the particle diffused into 2D AOI range (circle with radius of $1.96\sigma$, $\sigma$ being uncertainty of 2D Gaussian fit of individual AOIs) after being out of the range for a continuous 200-ms interval. Such biding dynamics indicate that the individual AOIs were visited 15.8 ± 9.9 times (Fig. S11a). Residence time was defined as the complete time interval between the time stamp of a visit event when the particle was first detected inside the AOI and the last time stamp the particle was detected inside the AOI. Residence time distribution was fitted into an exponential decay with one or more independent rate constants, as is shown in the following equation:

$P = \sum_{i=1}^{N} C_i e^{-k_i t}$ Here $N$ $(N = 1,2,\ldots)$ factors independently determine the frequency of duration of residence events in continuous 100-ms intervals. $C_i$ $(i = 1,2,\ldots,N)$ are prefactors and $k_i$ $(i = 1,2,\ldots,N)$ are rate constants. Half lives of biding were then can be obtained via $T_i = \frac{-ln2}{k_i}(i = 1,2,\ldots,N)$ , which gave a half life of 2.36 (95% confidence interval: 1.91, 3.12) second for single-factor binding and 0.28 (0.25, 0.32) and 5.96 (4.99, 7.42) second for bi-factor binding, respectively (Fig. S11b-c). It is noticeable that if diffusion is not limited on a 2D-surface, the average dwelling time $\tau$ of a particle undergoing Brownian motion should be:

$$\tau = \frac{\sigma^2}{2D}$$

Here $\sigma$ is the radius of a circular area of interest and $D$ is diffusion coefficient. $\sigma$ by our definition is 1.96 times of fitted AOI scale, which is 108 nm and $D$ is obtained from evaluation of 2-dimentional trajectory segments of 5s projected onto cylinder surfaces and was calculated to be 0.035 ± 0.011 μm$^2$/s. Taking these into account, the average duration of observing particle in the specified range is 0.18 s. This is comparable to the 0.28 s half-life of short-life events in the the bi-factor model fits and therefore indicates that such transient observation of particle visiting the restricted area could be non-binding freely diffusing events.

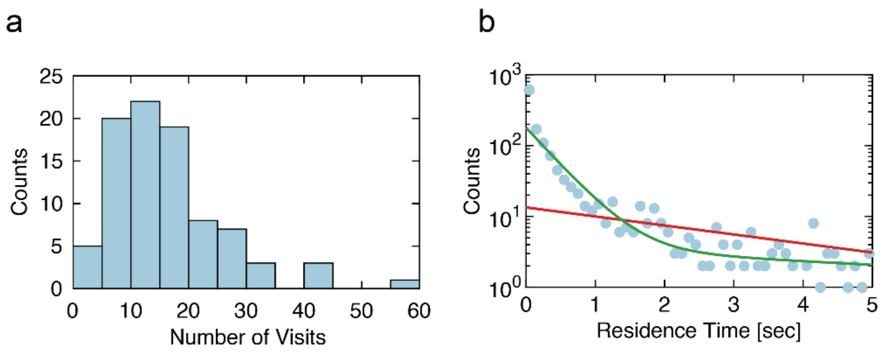

**Figure S11.** (a) Number of visits observed in each AOI. (b) Exponential fit of residence time from all visits in all labelled AOIs. Red curve and green curve indicate 1-factor and 2-factor fits, respectively.

**Supplementary Figure 12: Full image of trajectory of functionalized AgNP interacting with cell filopodia**

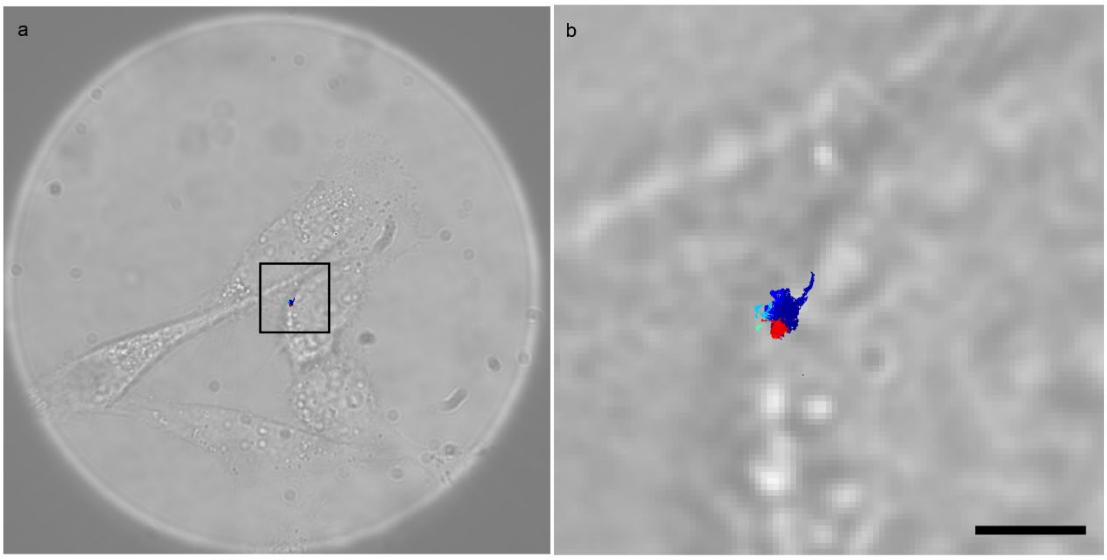

**Figure S12.** (a) Full trajectory shown in Figure 5 overlaid by bight field image immediately obtained after completion of the trajectory. Box dimension: 10 x 10 um (b) Closer view of box shown in (a) Scale bar: 2um.

### Supplementary Figure 13: Colocalization TATp functionalized AgNP and actin

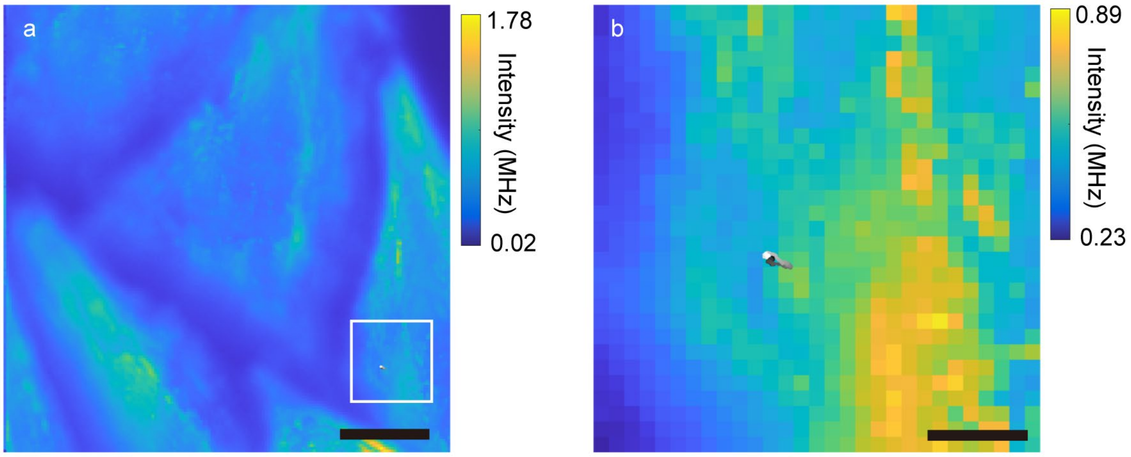

**Figure S13.** (a) TATp-Agnp trajectory (gray) with fluorescence image of actin (palura). The fluorescence image is a maximum intensity projection image of a 3D stack images. Scle bar: 5 μm. (b) Magnified view of (a). Scale bar: 2 μm.
